# Tactile stimulation transiently disrupts encoding of whisker position by cerebellar molecular layer interneuron ensembles

**DOI:** 10.1101/2025.08.12.669821

**Authors:** Elisabeth M M Meyer, Paul Chadderton

## Abstract

Molecular layer interneurons (MLIs) within the cerebellar cortex mediate feed-forward inhibition onto Purkinje cells, positioning them as pivotal modulators of cerebellar output. We asked how motor patterns and salient sensory input influence MLI population activity during active whisking and tactile interactions. Utilizing two-photon calcium imaging combined with high-speed videography, we examined MLI population dynamics in awake, behaving mice engaged in voluntary whisker movements. Our results demonstrate that during free whisking, MLI population activity reliably tracks whisker position, yielding uniform graded responses that provide stable and precise representations over time. Tactile contact with external stimuli evokes additional activation in a subset of MLIs. Sensory input transiently disrupts the linear relationship between MLI activity and whisker position and may account for rapid synchronous inhibition of Purkinje cell spiking activity following tactile stimulation. These findings indicate that MLIs in Crus 1 maintain an accurate internal model of whisker position which is perturbed upon encountering obstacles. This disruption likely reflects the integration of sensory feedback, disrupting predictive signals that are subsequently passed forward to Purkinje cells. Such signals can alert the brain to novel happenings and be used to update motor commands, thereby contributing to cerebellar function in response to environmental changes.

## INTRODUCTION

The brain facilitates interactions with the environment by anticipating future movements and their sensory consequences (Keller & Mrsic-Flogel, 2018). This task is achieved through internal representations that combine motor intentions and sensory feedback (Ito, 2012; Wolpert et al., 1998). The cerebellum is considered a key site for the formation of such representations and plays a critical role in planning and executing of movement (Tanaka et al., 2021), including control of the rodent whisker system (Rahmati et al., 2014). Rodents use their whiskers for tactile sensing, actively and rhythmically sweeping them to interact with their proximal surroundings. This behaviour is served by both efferent motor- and afferent sensory signals, which are received by the same regions of the cerebellum including lobule Crus1 (Proville et al., 2014). The principles by which the cerebellar cortex processes correspondent motor and sensory-related information is not well understood.

During voluntary movement, cerebellar Purkinje cells modulate their firing rates such that changes in simple spike rates correlate with changes in whisker position (Brown & Raman, 2018; Chen et al., 2016). Whisking in air, i.e. in the absence of external sensory input, can elicit both excitatory and inhibitory modulations to movement (Chen et al., 2016; Zhai et al., 2024). In contrast, tactile stimulation of the whisker pad evokes sharply timed decreases in Purkinje cell simple spike activity (Bosman et al., 2010; Brown et al., 2024; Brown & Raman, 2018). Rapid decreases in simple spike activity implicate a role for synaptic inhibition in Purkinje cell sensory responses via activation of molecular layer interneuron (MLIs). MLIs form inhibitory synapses onto Purkinje cell dendrites and provide feed-forward inhibition onto Purkinje cells (Eccles, 1967; Mittmann et al., 2005). Previously, MLIs have been shown to represent movement-related information (Astorga et al., 2017; Gaffield & Christie, 2017; Jelitai et al., 2016) in a similar manner to Purkinje cells (Chen et al., 2017). However, it is not clear how motor and sensory-related signals are organised at the level of MLI populations, and it is not known how MLI activity may contribute to Purkinje cell activity during active sensory processing.As MLIs provide the major source of inhibition to Purkinje cells, and inhibition of simple spiking appears to be the central means of representing tactile information in the cerebellar cortex, we asked how the MLI population represents motor and salient sensory stimulation information during free whisking and whisker touch, and whether, like Purkinje cells, MLIs employ a linear algorithm to represent whisker position. Using 2-photon imaging in the cerebellum of awake, freely whisking mice, we find that MLI ensembles in Crus 1 are active in a uniform, graded manner during free whisking. Tactile stimulation of the whisker produces additional activation within a smaller subgroup of MLIs. Using a linear model of MLI fluorescence changes, we were able to precisely reconstruct moment-to-moment whisker position from MLI population activity alone, showing that MLIs employ a linear algorithm for representing whisker position in real-time. This linear relationship between MLI population activity and whisker position is transiently disrupted during the touch periods, when tactile input causes additional MLI activation which degrades the correlation with whisker position. This additional MLI activation, superimposed on movement-driven signals, is appropriate for rapid silencing of downstream Purkinje cells during active touch.

## RESULTS

### Imaging the MLI population in Crus 1 reveals widespread graded responses to whisker movement

Crus 1 in the lateral cerebellar cortex processes sensorimotor information from the whiskers (Bosman et al., 2010; Bower & Woolston, 1983; Brown & Raman, 2018; Chadderton et al., 2004; Chen et al., 2016; Proville et al., 2014; Shambes et al., 1978). Most Crus 1 Purkinje cells display increases in simple spike activity during free whisking. In around one third of cells whisking onset produces reductions in tonic simple spike firing (Chen et al., 2016), supporting a role for inhibition from MLIs in cerebellar representation of voluntary movement. However, little is known about organisation of whisker movement signals in MLIs themselves. Therefore, we first sought to characterise how MLI ensembles respond during free whisking. To record the MLI population response to voluntary whisking, we utilized 2-photon imaging of Crus 1 in awake, head-fixed mice (**Fig. 1A**). By expressing GCaMP6f specifically in interneurons in the molecular layer (see Materials and Methods), we could selectively visualize these neurons and indirectly monitor their activity via changes in fluorescence ratio (dF/F) corresponding to intracellular calcium fluctuations. We simultaneously recorded whisker position during periods of active whisking and quiescence.

**FIGURE 1:**
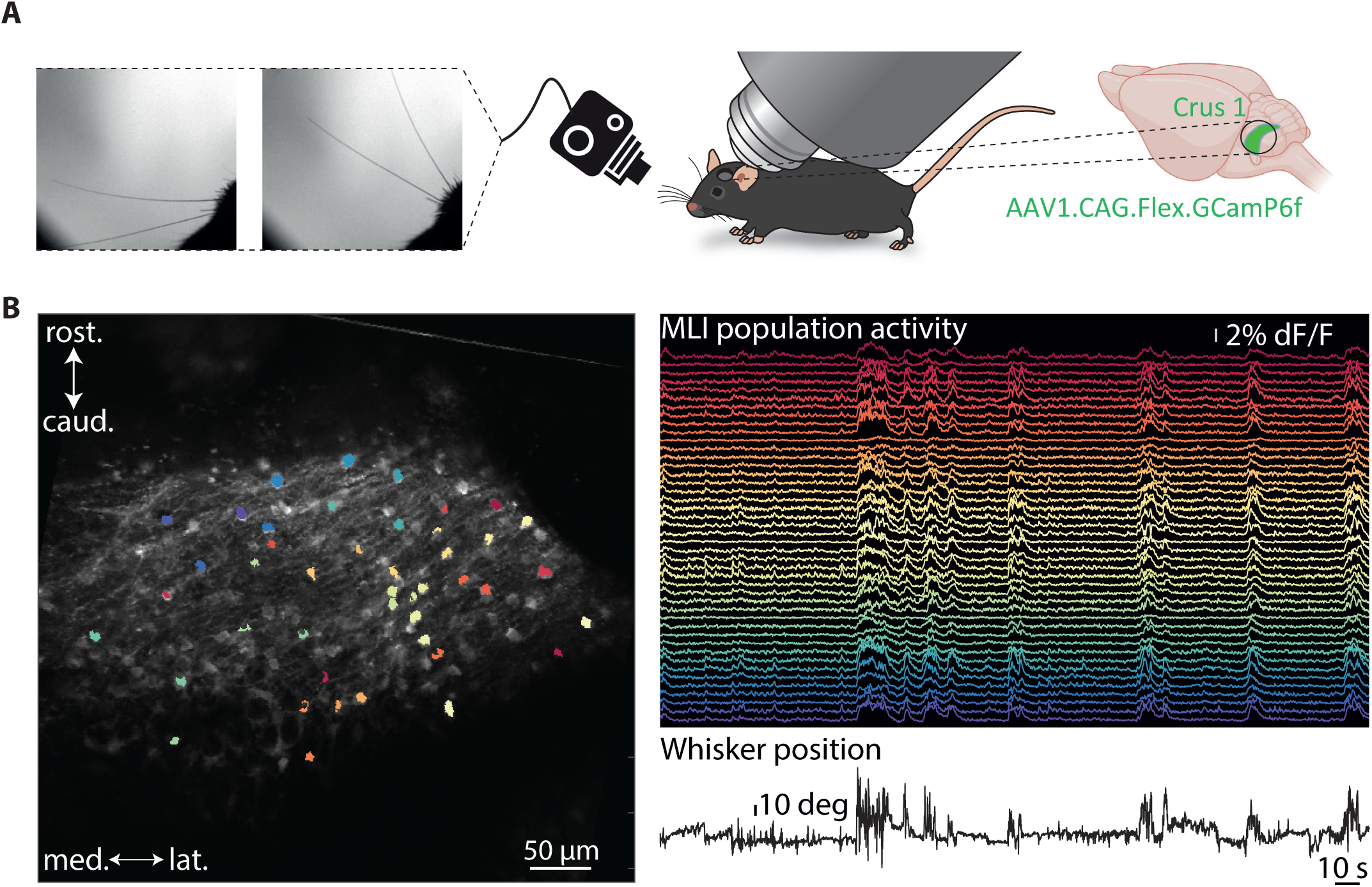
2-photon imaging of MLIs in Crus 1 during active whisking. A: Overview of experimental setup including position of the objective of the 2-photon microscope and high-speed camera for whisker recording. B: Whisker position and individual dF/F traces recorded from MLIs in awake, freely whisking mice and corresponding MLI cell body distribution throughout the FOV.

Calcium imaging revealed prominent increases in Crus 1 MLI activity during periods of whisking (**Fig. 1B**). We recorded 328 MLIs in 12 mice (27 ± 5 MLIs per animal). The population average of all recorded MLIs in Crus 1 showed a clear increase in activity upon whisking onset (*N* = 12 mice, *n* = 328 cells, average dF/F response amplitude: 0.021 ± 0.001, **Fig. 2A-C**). When aligned to movement onset, we observed that the MLI population responded in a graded manner, with individual MLIs varying in strength from silent to strongly responsive (**Fig. 2A**). Electrophysiological recordings previously revealed tuning to whisker position in single granule cells and inhibitory interneurons (Chen et al., 2017), and Purkinje cells (Brown et al., 2024; Brown & Raman, 2018; Chen et al., 2016). We therefore tested whether we could measure tuning to whisker position in fluorescence measurements from individual MLIs. On a cell-by-cell basis, we examined the relationship between dF/F and whisker position, to construct individual tuning curves (see Materials and Methods). Consistent with electrophysiological recordings, MLI fluorescence responses were well-tuned to whisker position. Within individual fields of view, MLIs showed broadly similar tuning profiles, exhibiting progressive increases in dF/F in both directions as positions deviated from the resting point (**Fig. 2D, E**). This contrasts with the response profiles of individual mossy fibres and granule cells (Chen et al., 2017), but is consistent with population firing rates measured via multisite microelectrodes (Palacios et al., 2025). In summary, we observe robust and salient increases in MLI activity in Crus 1 during voluntary whisker movements, supporting a role for MLIs in shaping simple spike firing responses in downstream Purkinje cells.

**FIGURE 2:**
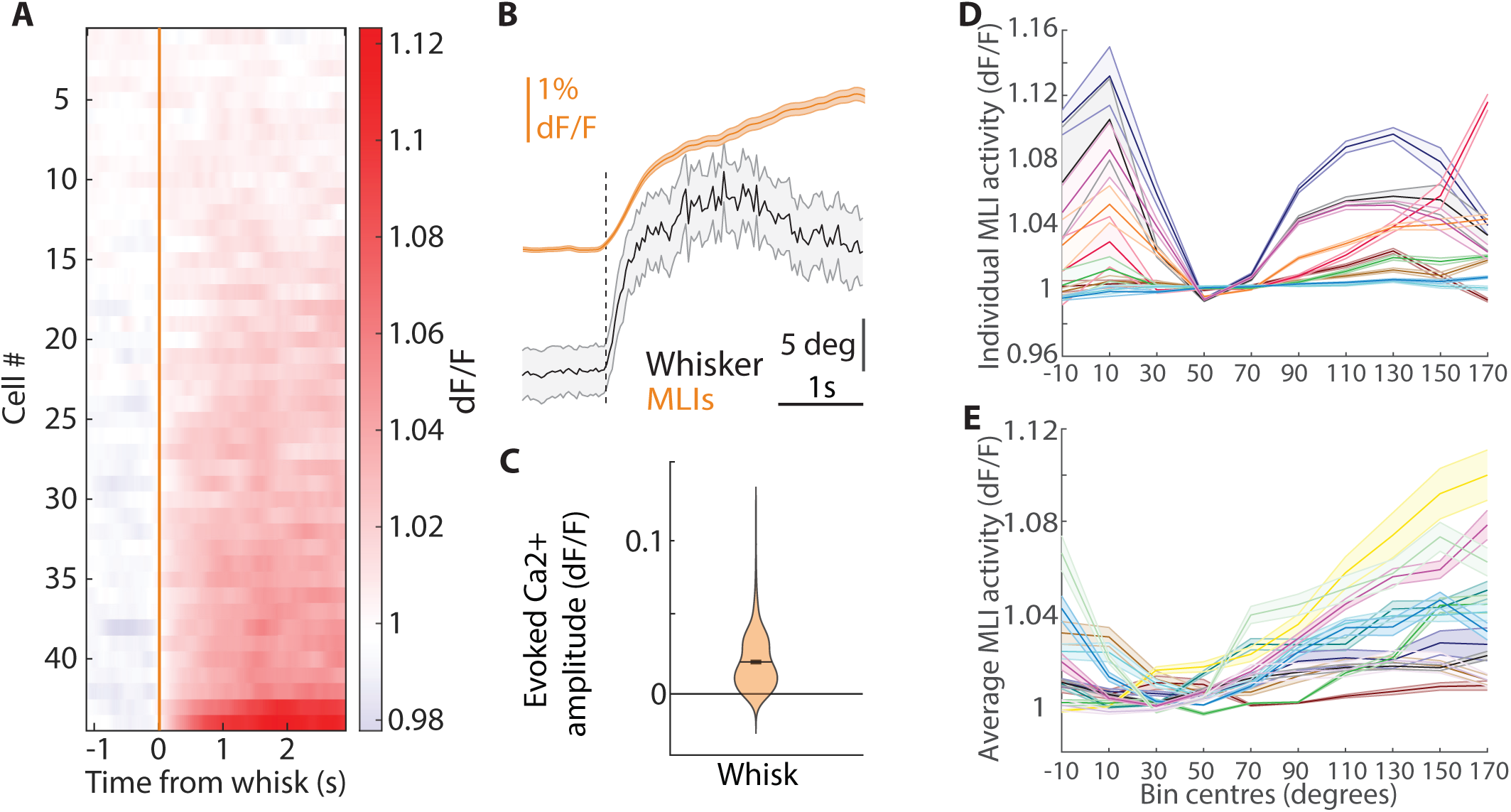
Whisking evokes graded uniform MLI activation in Crus 1. A: Evoked changes in Ca^2+^ (dF/F) in the MLI population of a single animal upon whisking (*n* = 44 neurons). B: Population response of all MLIs (*n* = 328 neurons) aligned to onset of whisking. C: Average amplitude of evoked change in Ca^2+^(dF/F) for all animals (*N* =12 animals, *n* = 328 neurons) aligned to whisking onset. D: Tuning curves of whisker position for 10 representative MLIs within a single field of view. E: Average tuning curves for multiple animals (*N* = 12 animals). Error bars = SEM.

### MLI population activity sufficient for whisker position prediction

Previous results have shown that spike trains from individual cerebellar interneurons are sufficient to modestly recover whisking trajectories during free whisking (Chen et al., 2017). Although imaged MLIs were tuned to whisker position, we were uncertain if the calcium-dependent fluorescence ratio dF/F could be similarly informative about whisking trajectories. To test this, we used a similar approach as has been previously used for spiking data (Hill et al., 2011). We averaged dF/F activity of each MLI population over a 20s window to calculate a linear transfer function (see Materials and Methods for details, **Fig. 3A**). We used this transfer function to predict whisker position during the remainder of the recording (4 minutes 40s; **Fig. 3B**). To assess the quality of the prediction, cross-correlation was computed between real and predicted movement patterns (**Fig. 3C**). As for electrophysiologically recorded spike trains, the linear transfer function was able to capture transformation dynamics of neuronal information to motor output faithfully over a long period. When the fluorescence signal was randomly shuffled in time, prediction quality degraded significantly as expected (*N* = 12 animals, aligned: 0.66 ± 0.073, shuffled: 0.29 ± 0.062, Wilcoxon signed rank test, *p* = 0.0024, **Fig. 3C**), indicating that whisker position could not be faithfully reconstructed. Taken together, these results show that the fluorescence signal from local MLI populations can be used to recover movement trajectories, and that underlying cerebellar representations of whisker movement remain stable over time. MLI population activity therefore maintains faithful representation of whisker movement, representing an internal model that can be applied to estimate current and future position of the whiskers.

**FIGURE 3:**
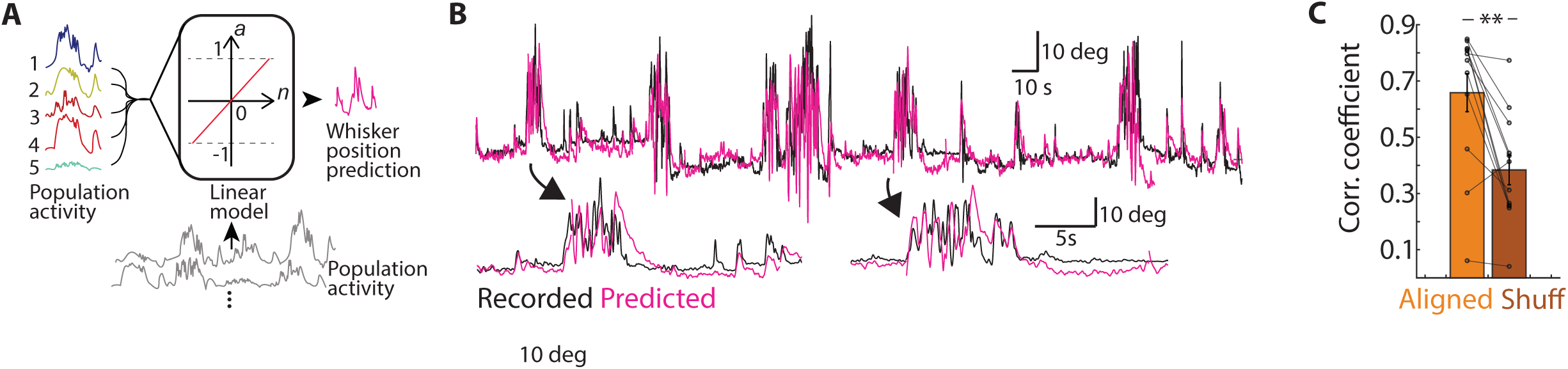
Reliable prediction of whisker position during self-generated motion from MLI population Ca^2+^ activity. A: Schematic of analysis approach. Linear transfer function predicts moment-to-moment whisker position from MLI population activity. B: Example period of recorded-(black trace) and reconstructed whisker position (pink trace) during free whisking for one animal. C: Correlation coefficients between measured whisker position and MLI-based reconstruction recorded within single fields of view (orange bar versus shuffled; brown; see Materials and Methods).

### Tactile stimuli elicit additional activation within Crus1 MLI populations

Whiskers not only provide rodents with information about their own position, but also transmit tactile information obtained through contact with objects in the environment. Amongst Purkinje cells, tactile whisker stimulation typically leads to rapid interruptions in simple spike firing that are highly synchronised across cell populations (Brown et al., 2024). We considered that inhibition of simple spiking could be driven by MLI populations responding to sensory-driven excitatory parallel inputs. In such a scheme, it is unclear how sensory-driven inputs might interact/interfere with widespread movement-related signalling. We therefore asked how MLI populations encode active touch. To do this, we made further MLI recordings after introducing a static pole close to the ipsilateral whisker pad. Animals were free to explore the pole using their whiskers and typically made frequent pole touches during individual recording sessions. Pole touches were identified using changes in whisker curvature (see Materials and Methods) (**Fig. 4A)**. Compared to movement-only trials (**Fig. 4Bi**), touch trials evoked stronger MLI activation (**Fig. 4Bii**). Touch events occurred at irregular times during individual whisking bouts so we separately aligned neural imaging data to touch onset. (**Fig. 4Biii**). When comparing data aligned to movement- and touch onset, we observed that MLIs display strong, graded responses at movement onset and additional activation following touch (**Fig. 4C**). Quantifying the peak changes in dF/F confirmed that the response amplitude of touch trials was significantly larger than the amplitude of free whisking alone (*N* = 12 animals, *n* = 328 neurons, whisking alone: 0.019 ± 0.002, whisking + touch: 0.029 ± 0.001, Wilcoxon signed rank test, *p* < 0.001, **Fig. 4D**). These data demonstrate that individual MLIs multiplex touch-evoked and self-generated signals. By combining both movement and touch-related information, MLIs can provide elevated levels of inhibition to Purkinje cells during tactile stimulation.

**FIGURE 4:**
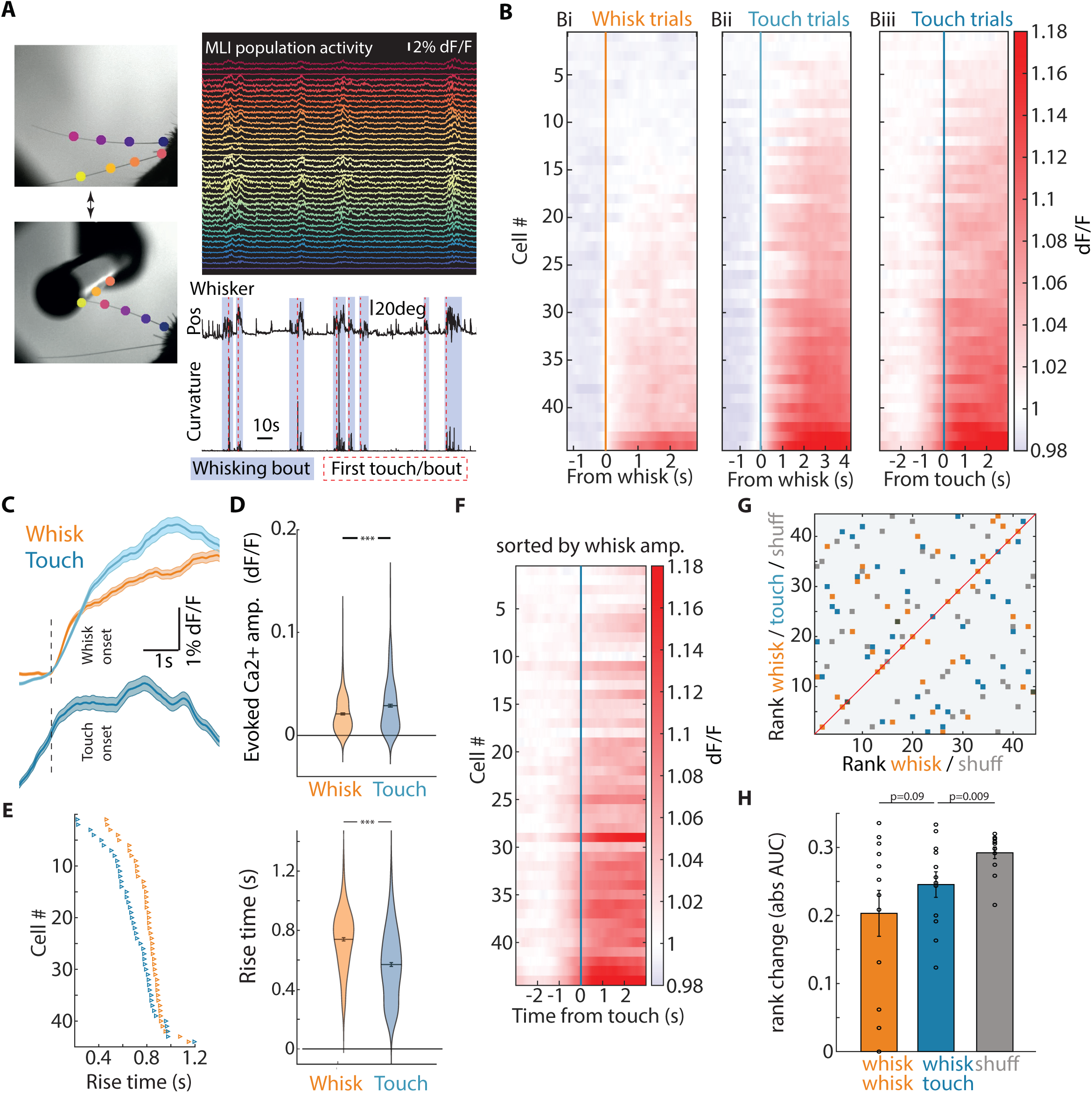
Tactile stimulation evokes additional Ca^2+^ activity in a subset of Crus 1 MLIs. A: Left column: Video stills of the whiskers during free whisking (top) and when the animal is touching the pole (bottom) (dots: DeepLabCut tracking markers). Right column: Example dF/F activity during pole touching (top), whisker position (Pos) with whisking bouts marked in light blue and changes in whisker curvature with red dotted lines marking the first pole touch for each whisking bout (bottom). Same animal as in Figure 2. B: Example MLI population response aligned to whisking onset for whisking only (Bi) and touch trials (Bii). Biii, touch trials aligned to touch onset. All plots sorted by response amplitude. Same animal as in Figure 2. C: Top, Average response of all MLIs recorded for whisking only (orange), and touch trials (light blue), Bottom, average response of touch trials aligned to touch onset (dark blue). D: Peak MLI Ca^2+^ activity during free whisking trials (orange) and touch (blue). E: left, Ca^2+^ event rise times for free whisking - (orange) and touch trials (blue) for the same animal as in B. Right, average rise times across all animals (*N* = 12 animals) for free whisking (orange) and touch (blue). F: Touch responses sorted by size of whisking responses for one animal (same animal as in B). G: Change in rank for each MLI for the animal in F for 3 different conditions: free whisking versus touch (blue), free whisking versus free whisking (orange) and control (free whisking versus arbitrarily shuffled dataset control; grey). H: Quantification of the area under the curve (AUC) as a measure of changes in rank (see Material and Methods) and thereby changes in response size between different conditions. Error bars = SEM.

Purkinje cell population activity following sensory stimulation has been shown to be highly synchronized, resulting in rapid and concerted decreases in simple spike activity in response to tactile stimuli (Brown & Raman, 2018). Importantly, such changes in simple spike activity are faster than movement-related modulations. Therefore, we asked whether touch-evoked activation was more rapid within MLI populations. To do this, we compared MLI activity rise times (see Materials and Methods) between whisking bouts and touch events. We found that rise time of activity across the population was significantly faster for touch events than for whisking onsets (*N* = 12 animals, *n* = 328 cells, whisk alone: 0.74 ± 0.011s, touch: 0.57 ± 0.013, Wilcoxon signed rank test, *p* <0.001, **Fig. 4E**). This shows that MLI activity is sharply timed during tactile stimulation, whereas efferent information is transmitted over broader time scales, supporting the proposal that MLIs contribute significantly to shaping temporal response characteristics of Purkinje cell during both afferent and efferent bouts of activity.

Previously we observed a gradient of activation with local MLI populations. We therefore asked whether MLIs responding strongly to whisker movement alone also responded most strongly to touch, or, if different MLI ensembles preferentially represent each condition. We ranked MLIs according to the peak amplitude of their response during whisker movement alone and found this ranking to be a poor predictor of the MLI response to touch (**Fig. 4F**). We then compared MLI rankings based on response amplitude during movement-only and touch bouts (**Fig. 4G**). Overall, while some neurons retain their rank within the population (17 ± 1%), thereby exhibiting similar response strength to touch as to whisking, most MLIs do not exhibit a predictable touch response based upon their whisking response (larger touch response than whisking response: 40 ± 1%, smaller touch than whisking response: 43 ± 1%).

When plotting the change in rank between movement-only and touch bouts, computing the AUC for the resulting curve and comparing it to a shuffled dataset and a within-condition (whisk-whisk), we found differences in absolute AUC for the changes in rank between whisking and touch, for the randomly shuffled dataset and the within-condition changes in rank (*N* = 12 animals, whisk-touch: 0.25 ± 0.018, shuffled: 0.29 ± 0.01, whisk-whisk: 0.20 ± 0.034, Wilcoxon signed rank test, whisk-touch vs. shuffled: *p* = 0.0093, whisk-touch vs. whisk-whisk: *p* = 0.092, **Fig. 4H**). This indicates that the level of MLI activation during bouts of voluntary whisking does not predict how the same cells will respond to active touch. These findings together suggest that MLIs might form dedicated separate ensembles to encode motor signals and active touch. Ensembles could be formed by groups of MLIs receiving biased parallel fibre input conveying mainly either self-generated sensorimotor information or tactile stimulation, thereby generating MLI ensembles with different response profiles.

### Tactile signals transiently disrupt whisker position predictions

During free whisking, MLI populations can precisely represent whisker position in a stable manner over long time scales using a linear-based reconstruction (**Fig. 3**). We questioned whether a linear reconstruction based on neural imaging data would further be able to recover whisker position during tactile stimuli events. As before, we used data obtained during the first 20 seconds of the imaging session to calculate the linear transfer function during a period of free whisking without touches. We then applied the transfer function to the remainder of the neural imaging data to predict whisker position, including during periods of active touch. We found that the linear relationship between MLI activation and whisker position transiently breaks down during active touch (**Fig. 5A, B**), (*N* = 12 animals, whisking alone: 0.36 ± 0.07, touch: 0.13 ± 0.05, Wilcoxon signed rank test, *p* = 0.043, **Fig. 5C**). Additional activation caused by tactile feedback from the pole touch disturbed the relationship between MLI population activity and whisker, reducing the instantaneous correlation between predicted and measured whisker position compared to voluntary whisking. Movement prediction signals in the cerebellar molecular layer are thus transiently disrupted at moments of tactile sensory input.

**FIGURE 5:**
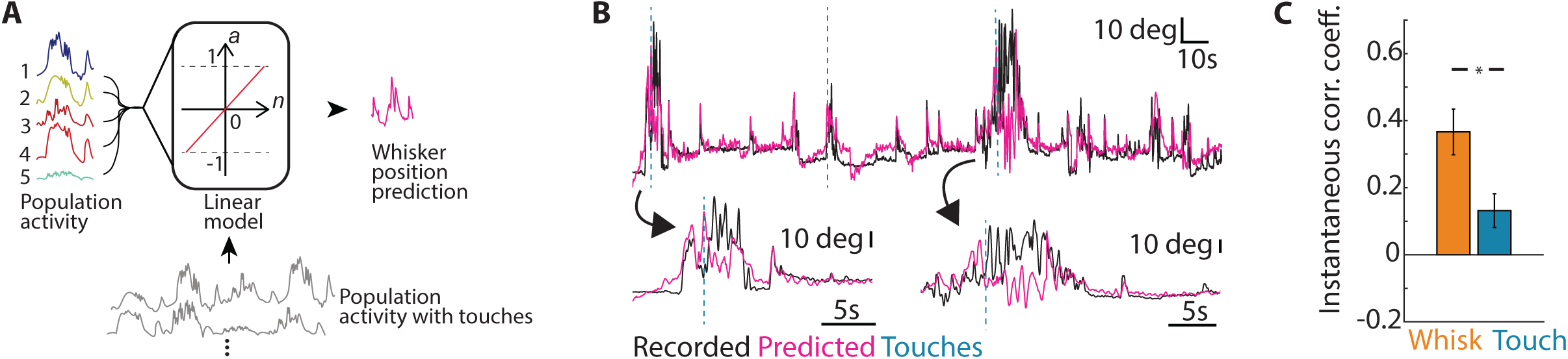
MLI prediction of whisker position breaks down transiently during tactile stimulation. A: Schematic of analysis approach. A linear transfer function is calculated from 20s of whisking only data (no touch). Predictions are subsequently tested on data which includes pole touches. B: Example recorded (black trace) and reconstructed whisker position (pink trace) during pole touching for one animal (same animal as in Figure 3). C: Instantaneous correlation between recorded and reconstructed whisker position during whisking only (whisk, 1s window at whisking onset) and around the time of pole touching (touch, 1s window at pole touch).

## DISCUSSION

By applying 2-photon imaging to the cerebellum of awake, freely whisking mice, we show that MLI ensembles in Crus 1 exhibit a consistent, graded activation pattern during whisking. Movement-related activity is distributed in a widespread and coherent manner (Gaffield & Christie, 2017), while a subset of MLIs are additionally engaged by tactile sensory input. Using a linear model, we accurately reconstructed the whiskers’ moment-to-moment positions based solely on MLI population activity, demonstrating that MLIs use a linear coding strategy to represent whisker position in real-time. This linear relationship between MLI activity and whisker position was disrupted during touch events, indicating that tactile input transiently interferes with the cerebellar representation of whisker position.

MLIs provide feed-forward inhibition to Purkinje cells in response to motor and sensory signals, and are therefore positioned to exert significant control over Purkinje cell firing and cerebellar cortical output (Eccles, 1967). Direct electrophysiological recordings have previously shown that MLIs are tonically active when the animal is at rest (Armstrong & Rawson, 1979; Jorntell & Ekerot, 2003), and can both increase or decrease their firing rate upon engaging motor behaviour such as whisking (Chen et al., 2017). However, indirect activity measures such as calcium imaging have consistently reported uniform activation of MLIs upon behavioural engagement (Astorga et al., 2017; Gaffield & Christie, 2017). Here we also observed uniform increases in MLI activity (though graded in amplitude) within local populations during whisking. We did not observe MLIs that significantly decreased their activity during whisking. Previously observed reductions in MLI firing rate have been in the range of a ∼10Hz decrease from baseline (Chen et al., 2017). Such small reductions in firing rate might be impossible to resolve using GCaMP6f, and it not known how the reporter interacts with endogenous calcium buffering processes within MLIs. We hypothesize that minimally activated MLIs in our data could be inhibited populations observed via electrophysiological methods (**Figs. 2A, 4B**).

Recent work has shown that MLIs can be separated into two subpopulations based on their postsynaptic targets and effect on Purkinje cells; it was however found that the majority of MLIs directly inhibit Purkinje cells (Lackey et al., 2024), with a smaller number providing disynaptic disinhibition. This suggests that our MLIs recorded during both whisking and touch are dominated by cells which exert direct inhibitory effects on Purkinje cells. We found that rise times of the Ca^2+^ signal in MLIs was significantly shorter for touch events compared to whisking only (**Fig. 4E**), implying an underlying increase in spiking that is more tightly constrained in time during tactile sensation compared to movement alone. Purkinje cell touch responses have been characterized by synchronous, well-timed inhibition of simple spiking (Brown et al., 2024), while movement responses in Purkinje cells can be both excitatory and inhibitory (Chen et al., 2016), gradual changes in firing rate (Brown et al., 2024). Together this suggests that MLIs transmit sharply timed information during tactile whisker stimulation, whereas efferent information is transmitted over broader time scales. The difference in rise times could lead to differences in how excitation from parallel fibres and inhibition from MLIs interact in Purkinje dendrites, with sharp inhibition leading to synchronized breaks in Purkinje cell firing, while slower rising combinations with parallel fibre excitation arriving at different time scales at the Purkinje cell dendrite leading to more complex excitation/inhibition interactions and perhaps explaining the variability of Purkinje cell responses to movement initiation (Hong et al., 2016).

By tracking the activity of identified MLIs during both movement and sensory stimulation, we showed that it is not the same MLIs that are strongly excited by both whisking and touching, i.e., MLIs strongly responding to whisking (movement only) are not necessarily be strong responders to touch (sensory stimulation), and vice versa. This suggests that MLIs might form functional subpopulations. These functional ensembles may be targeted by different groups of parallel fibres carrying sensory-versus efferent information and may further target different Purkinje cells or exert different postsynaptic effects. Indeed, during activation by naturalistic stimuli, it has been shown that neighbouring MLIs and Purkinje cells are unlikely to be driven by the same granule cells and thereby would be receiving different parallel fibre inputs (Jorntell et al., 2010). Considering that Purkinje cells similarly form functional ensembles that exhibit specialized activity patterns which are associated with specific phases of movement or sensory input (Chang et al., 2020; Herzfeld et al., 2015; Hoogland et al., 2015), functional MLI ensembles could be organized into complex microcircuits that generate motor-or sensory-specific inhibition and form processing clusters dedicated to either sensory or efferent encoding with subpopulations of Purkinje cells. Cerebellar network activity has been shown to encode both predictions of motor command consequences and corresponding sensory feedback of these motor commands (Popa et al., 2013; Streng et al., 2018; Wolpert et al., 1995), with a bias towards motor predictions over sensory feedback (Hewitt et al., 2011). Ongoing motion can be controlled and adjusted based on a mismatch between motor predictions and sensory feedback, which is described as a prediction error (Popa & Ebner, 2018). Our data show that during voluntary free whisking, there is a robust linear relationship between MLI activity and whisker position, as shown through reconstruction of moment-to-moment whisker position from neural data alone (**Fig. 3**). This linear relationship is transiently interrupted by tactile stimuli (**Fig. 5**). This suggests that touch events act to disrupt the expectation of being able to freely move whiskers through space, essentially generating a prediction error which is forwarded to Purkinje cells. This inhibitory prediction error could be used by the cerebellum to signal that its internal model of the world is unreliable (Palacios et al., 2021; Tanaka et al., 2021). Inhibition plays important roles in predictive coding by suppressing expected inputs, thereby making coding sparser and more efficient, and maintaining the dynamic range of the network and preventing run-away excitation (Keller & Mrsic-Flogel, 2018). Considering the inhibitory nature of Purkinje cells, our data imply inhibition by MLIs acts not just as suppressor, but also as an actuator (Heiney et al., 2014). Suppressing Purkinje cells during unexpected stimuli leads to disinhibition in the deep cerebellar nuclei and increased cerebellar output to downstream targets signalling the unexpected tactile input (Brown et al., 2024). Taken together, our data show that MLIs play a critical role in adaptive sensorimotor processing and support the cerebellum in generating accurate and flexible predictions based on constantly updated models of the external world.

## MATERIALS AND METHODS

The care and experimental manipulation of animals was performed in accordance with institutional and United Kingdom Home Office guidelines.

### Surgical procedures

Data was collected from a total of 12 8–12 week-old male and female nNos-Cre mice (B6.129-*Nos1^tm1(cre)Mgmj^*/J, #017526, The Jackson Laboratory). Virus injections into Crus 1 and cranial window implantation were performed as previously described (Dombeck et al., 2007; Leinweber et al., 2014). Briefly, mice were anaesthetized using Isoflurane (3-5% induction, 1.5% maintenance). Carprofen (5 mg/kg) and Buprenorphine (0.075 mg/kg) were injected intraperitoneally. Lidocaine (2mg/kg) was injected locally on the scalp and muscles attached to the occipital ridge (m. semispinalis capitis). The skin above the skull was removed and the m. semispinalis capitis gently removed to expose the location of the craniotomy over Crus 1. A ∼3mm craniotomy was drilled over left Crus 1 in the cerebellum. 5 viral injections (virus: AAV(1)-CAG-Flex-GCaMP6f, 5×10^12^ GC/ml, Lot #100835, Addgene) were performed at ∼200 μm depth at various locations within Crus 1 (injection volume: ∼250 nl per site), and a 3 mm diameter glass coverslip was fitted over the craniotomy. The exposed skull and muscle tissue was covered in Histoacryl (Braun), a custom designed headplate was fixed to the top of the skull (redesigned version of headplate shown in Leinweber et al., 2014) using dental cement (Paladur). Animals were continuously group-housed during surgery recovery (>21 days) and imaging. 3-4 days before imaging, all but 3 whiskers were trimmed on the left whisker pad (ipsilateral to the craniotomy).

### Imaging

For functional calcium imaging, animals were head-fixed on a custom-build stage under the 2-photon microscope (Hyperscope, Scientifica) equipped with 8 kHz resonance scanner, which enabled recording speeds of 31 Hz for 512×512 pixel frames. The angle of the objective (16x, NA 0.80, Nikon) was set to 30 degrees for all experiments (vertical: 0 degrees). GCaMP6f was visualized using a femtosecond laser (Chameleon Ti:Sapphire, Coherent) tuned to 920 nm. During the experiment, animals sat in a plastic tube and were free to whisk. A high-speed camera (DALSA GENIE-HM640) controlled via Streampix (Norpix) and an infrared LED were used to visualize and record whisking activity at 300 Hz ipsilaterally to the craniotomy. For touch experiments, a pole was intermittently introduced within the ipsilateral whiskers’ reach (alternating pole present/pole absent recording phases). For each animal, 15 mins of undisturbed voluntary whisking and 15 mins of pole touching was recorded.

### Analysis of imaging data

Raw images were registered using single-line motion correction implemented in Sima (Image processing software)(Kaifosh et al., 2014) and full-frame registration in Calliope (Image processing software, https://svn.code.sf.net/p/iris-scanning/calliope/) to correct for brain-motion artifacts in the x-y plane. Datasets that still had motion artifacts after running both motion correction algorithms were discarded. MLIs were manually selected for calcium signal extraction based on the mean and maximum fluorescence projection throughout the whole experiment (Keller et al., 2012); only MLIs visible throughout the whole recording session were included in the analysis. Slow drift of raw fluorescence was corrected using an 8^th^ percentile filtering with a 15s sliding window (Dombeck et al., 2007) and dF/F was calculated as the mean fluorescence of each MLI in the frame normalized by the median of the fluorescence distribution. dF/F was further filtered using a 1^st^ order Savitzky-Golay filter with frame length 11 (Matlab, Mathworks). Further analysis was performed using custom-written scripts in Matlab.

### Whisker tracking

Individual whiskers were tracked using DeepLabCut (ResNet 50) (Mathis et al., 2018; Nath et al., 2019). 4 markers were placed on each whisker, the coordinates of which were fitted with a 4^th^ order polynomial to reconstruct the whisker. The whisker angle was extracted by fitting a tangent to the base of the reconstructed whisker and calculating its angle with respect to a vertical fitted through the centre of the nose of the animal. For further analysis, whisker angle data was down sampled to 31 Hz to match neuronal data.

Whisker curvature for touch detection was calculated by fitting a circle through the 3 most distant whisker markers (excluding the marker at the base of the whisker) and calculating its radius (Knutsen et al., 2008). This method does not identify putative ‘touch events’ where the whisker is not applied with enough force to bend it.

### Data analysis

Episodes of whisking activity were identified semi-automatically by transforming the whisker trace into a binary step function, extracting the steps as ‘whisking epochs’ and then manually optimising start and end points. Timestamps of whisking epoch starts were used to align dF/F activity for each neuron. Population averages were calculated by first averaging each neuron’s activity and then averaging over the whole population. Whisking-evoked amplitudes were calculated by averaging dF/F over 1.5 seconds (45 frames) from the whisking onset timestamp and subtracting 1s (31 frames) baseline activity directly preceding the respective whisking epoch. dF/F rise times were calculated as the time the calcium signal takes to cross from 10% to 90% of the maximum amplitude reached after the beginning of a whisking epoch and within 1.5 s (45 frames). For ranking neurons according to their response sizes, we quantified response amplitudes to whisking onset and touch onset as described above and sorted them according to their size, which gave each neuron a rank for whisking (only trials without pole touches) and a rank for touch. As a control, we separated half of the whisk-only trials and ranked neurons in either half, to get a sense of baseline trial-to-trial variability. Additionally, we randomly shuffled both whisking only and touch ranks for the ‘shuffled’ control. To quantify differences between ranks, we subtracted touch rank from the whisk rank/ whisk rank from the whisk rank respectively, and plotted the resulting delta rank in ascending order. We then quantified the absolute area under the curve of the resulting trace for each condition (whisk-whisk, whisk-touch and shuffled control).

To calculate tuning for individual neurons, we sorted dF/F values according to the corresponding whisk angle and then binned the resulting data by averaging over each 20 degree bin. To generate a tuning curve for each animal, we averaged all individual tuning curves.

To reconstruct whisking activity from neuronal activity, we averaged neuronal activity for each condition for each animal (N = 12, voluntary whisking, ∼ 15 mins recording time; pole: ∼ 15 mins recording time total). We used the first 20 s of the recording to estimate the transfer function using the tfest function for discrete time data in the System Identification Toolbox in Matlab (Mathworks). Briefly, we created an ‘iddata’ object containing the training data with the neuronal data as the system input and the whisker position as system output. Using this data object, we estimated a 2-pole, 2-zero linear transfer function, which we then used to reconstruct 20 s windows of the remaining recording time (14 mins 40 s) using the ‘compare’ function in the system Identification Toolbox in Matlab. Next, we concatenated the predicted responses and cross correlated the estimated whisker position with the actual recorded whisker position for the whole recording. As a control, we randomly shuffled the datapoints in the dF/F signal and used the already trained model to attempt to predict whisker position from the shuffled neuronal activity. For instantaneous predictions during whisking and touch (**Fig. 5**), we calculated the cross correlation between the recorded and the predicted whisker angle within 1s (31 frames) of a touch occurring, and compared this to the correlation within 1 s (31 frames) of a touch-free whisking epoch.

### Statistics

Values reported are mean and standard error of the mean (SEM), unless otherwise stated. All statistical tests performed are non-parametric. We used the 2-sided Wilcoxon signed rank test to test for statistical significance between groups unless otherwise stated.

## AUTHOR CONTRIBUTIONS

EMMM and PC designed the experiments, interpreted the data, and wrote the manuscript. All experiments were performed and analysed by EMMM.

## ACKNOWLEDGEMENTS

This work was supported by a Wellcome Trust Investigator Award (209453/Z/17/Z) to PC. The authors declare no competing interests.

